# Age-related Impairment of Implant Osseointegration is Associated with Immune Activation and Suppression of Angiogenic, Notch, and Wnt Pathways

**DOI:** 10.1101/2020.12.27.424462

**Authors:** Kathleen Turajane, Gang Ji, Yurii Chinenov, Max Chao, Ugur Ayturk, Matthew B. Greenblatt, Lionel B. Ivashkiv, Mathias PG. Bostrom, Xu Yang

**Affiliations:** Hospital for Special Surgery, New York, NY; The Third Hospital of Hebei Medical University, Shijiazhuang, China; Department of Pathology and Laboratory Medicine, Weill Cornell Medicine, New York, NY

**Keywords:** osseointegration, aging

## Abstract

The number of total joint replacements (TJRs) in the United States is increasing annually. Cementless implants are intended to improve upon traditional cemented implants by allowing bone growth directly on the surface to improve implant longevity. One major complication of TJR is implant loosening, which is related to deficient osseointegration in cementless TJRs. Although poor osseointegration in aged patients is typically attributed to decreased basal bone mass, little is known about the molecular pathways that compromise the growth of bone onto porous titanium implants. To identify the pathways important for osseointegration that are compromised by aging, we developed an approach for transcriptomic profiling of peri-implant tissue in young and aged mice using our murine model of osseointegration. Based on previous findings of changes of bone quality associated with aging, we hypothesized that aged mice have impaired activation of bone anabolic pathways at the bone-implant interface. We found that pathways most significantly downregulated in aged mice relative to young mice are related to angiogenic, Notch and Wnt signaling. Downregulation of these pathways is associated with markedly increased expression of inflammatory and immune genes at the bone-implant interface in aged mice. These results identify osseointegration pathways affected by aging and suggest that an increased inflammatory response in aged mice may compromise peri-implant bone healing. Targeting the Notch and Wnt pathways, promoting angiogenesis, or modulating the immune response at the peri-implant site may enhance osseointegration and improve the outcome of joint replacement in older patients.

## INTRODUCTION

The number of total joint replacements (TJRs) in the United States is increasing annually, including increased utilization in elderly patients well into the eighth or even ninth decade of life ^(1)^. Cementless implants are intended to overcome the shortcomings of traditional cemented implants by allowing bone growth directly onto their surface to improve implant longevity ^(2,3)^. Although most patients have good long-term outcomes, a major complication of TJR is implant loosening ^(4,5)^, which in cementless TJRs is related to deficient osseointegration, the physical and functional connection between bone and implant ^(6,7)^. One of the most important risk factors for poor osseointegration, and thus decreased implant longevity, is elder age of patients ^(8)^. Poor osseointegration in aged patients is typically attributed to decreased basal bone mass ^(9,10)^. Little is understood about the mechanisms which compromise de novo bone formation – bone ingrowth into porous titanium implants in the elderly. To develop therapeutic strategies to improve clinical outcome of TJRs in elderly patients, we need to understand how aging compromises the molecular pathways important for osseointegration.

TJR surgery involves creation of a bone injury by drilling a canal into bone, followed by implant insertion and subsequent healing and ingrowth of cancellous bone into the porous surface of the titanium implant ^(11)^. We and others have developed animal models to analyze the process of orthopaedic implant integration ^(12–24)^. In our mouse model, load-bearing titanium implants inserted into the proximal tibia become integrated with the surrounding bone over a 4-week period ^(15,19,23)^. Little is known about molecular pathways that regulate this process. In other bone injuries such as fracture, repair initiates with a hematoma and an inflammatory phase followed by a bone formation phase whose mechanism (endochondral versus intramembranous) is determined by the extent of mechanical stability at the injury site. The determinants of effective bone repair include the nature, magnitude and kinetics of the inflammatory response, effective neovascularization, the functionality and mobilization of skeletal progenitor cells and osteoclasts, and induction of the canonical Wnt, BMP and Notch anabolic pathways ^(25,26)^. Little is known about the relative importance of these determinants in implant osseointegration.

Cross-regulation between the immune system and bone metabolism has long been appreciated and its study has been termed ‘osteoimmunology’ ^(27,28)^. Sustained high-level inflammation associated with infection, autoimmune diseases and foreign body reactions induces pathological bone resorption. Important mechanisms include augmentation of osteoclastogenesis directly by macrophage-derived inflammatory cytokines such as IL-1 and TNF, and indirectly by Th17 T cells via activation of stromal cells to express RANKL ^(29,30)^. Th17 cells are expanded and promote bone resorption in response to continuous elevation of PTH ^(31)^. Under inflammatory conditions, cytokines such as TNF and IFN-γ additionally suppress bone formation by inhibiting the differentiation and function of osteoblasts ^(32)^.

Immune-bone crosstalk is also important in the absence of overt inflammatory disease, and can play important roles in bone homeostasis, response to injury, and subsequent repair ^(27,28)^. An early, transient, low-level inflammatory response after bone fracture is mediated by inflammatory cytokines including TNF and IL-17 and is important for removal of dead cells and tissue debris, and priming of subsequent neovascularization and mobilization of PDGFRα+Sca1+ skeletal progenitors ^(33,34)^. The later phases of bone healing are promoted by trophic (also termed M2) macrophages that produce growth factors, and suppressed by IFN-γ-producing CD8+ T cells ^(35–38)^. Additionally, bone healing and repair can be promoted by regulatory T cells (Tregs) that suppress pathogenic functions of CD8+ T cells, and instead induce these cells to produce Wnt10b in the bone marrow ^(36,39)^. The overall model is that a balanced inflammatory response that evolves over time from type 1 inflammatory to type 2 pro-resolution immunity can promote bone healing, whereas excessive or unbalanced immune responses will delay and compromise repair.

Here, we sought to identify pathways important for osseointegration that are compromised by aging. In this study, we developed an approach to perform transcriptomic profiling of peri-implant tissue in young and aged mice using our murine model of osseointegration ^(23)^. We hypothesized that aged mice have impaired activation of bone anabolic pathways at the bone-implant interface. We found that pathways most significantly downregulated in aged relative to young mice are related to angiogenesis, Notch and Wnt signaling. This downregulation of bone formation pathways was associated with strikingly increased expression of inflammatory and immune genes at the implant-bone interface in aged mice. These results identify osseointegration pathways affected by aging, and suggest that an increased inflammatory response in aged mice may compromise bone healing.

## MATERIALS AND METHODS

### Study Design

Using an IACUC-approved protocol, 16-week-old (young) and 91-week-old (aged) female C57BL/6 mice (n=10 per group) were purchased from Jackson Laboratory (Bar Harbor, Maine). A porous titanium (Ti6Al4V) implant (Eosint M 270; Eos Electro Optical Systems, Munich, Germany) was inserted into the right proximal tibia of the mice as previously described (Figure 1) ^(15,19,23)^. Mice started weight-bearing with the implanted knee immediately after recovering from anesthesia.

**Figure 1.**
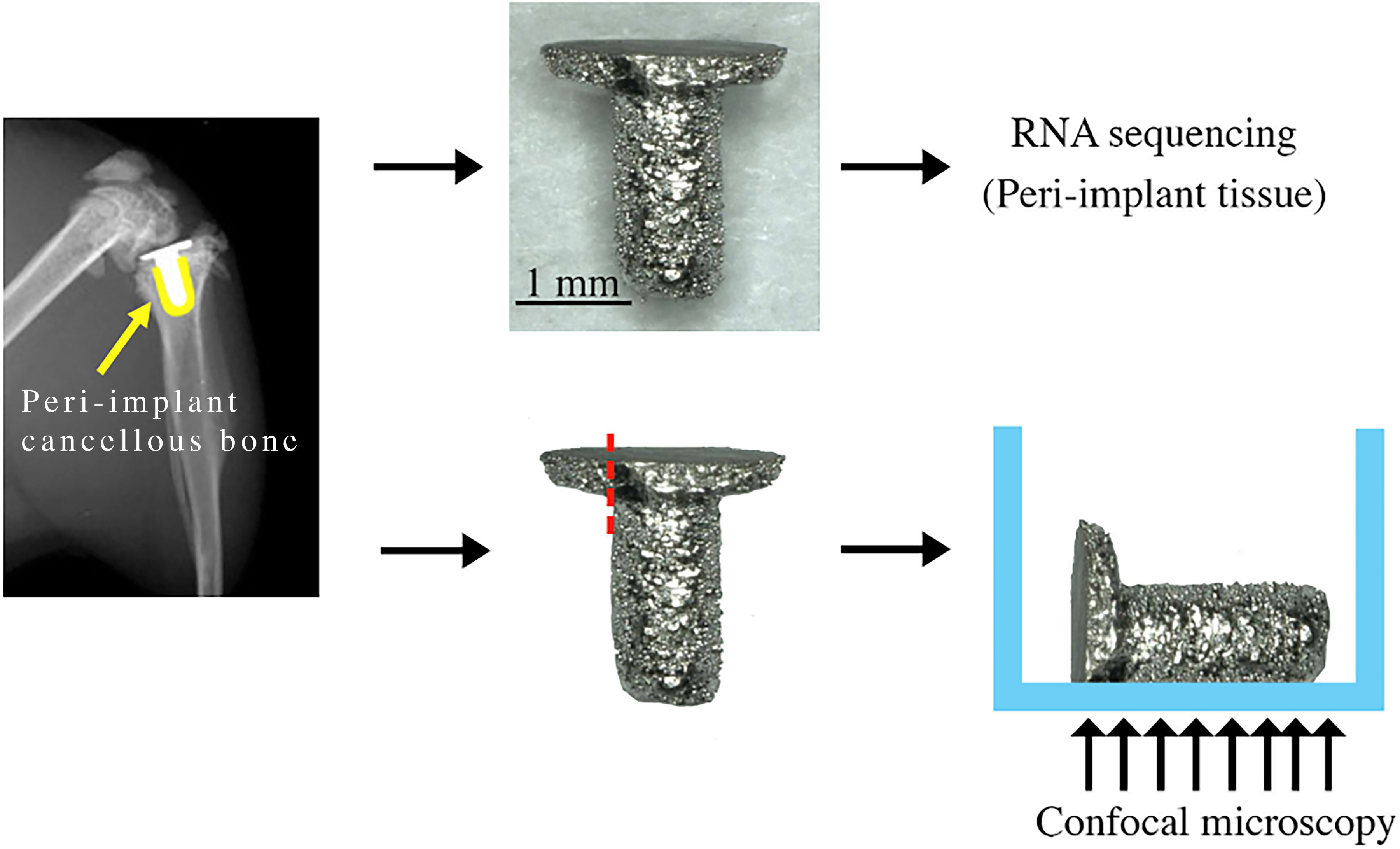
Peri-implant tissue collection and immunofluorescence staining preparation. Peri-implant tissue collection process. The implant was removed from the right tibiae. (Yellow: cancellous bone around the implant). One end of the tibial plateau component of the implant was polished before immunofluorescence staining.

### RNA extraction and purification

Mice were sacrificed one-week post-implantation. Right tibiae were rapidly dissected to remove soft tissue, fibulae, and malleoli. Implants with attached tissue (peri-implant tissue) were removed from the tibia and bone marrow was isolated by centrifugation (13,000 rpm for 30 seconds) using nested tubes. Next, the implant with tissue on its surface and adjacent cancellous bone, collected using a 1mm biopsy punch, were separately stored in 2 mL pre-filled 2.4mm metal beads tubes (VWR International, Radnor, PA), and quickly snap-frozen in liquid nitrogen ^(40)^. The bone specimens were stored at −80°C for later RNA extraction and purification.

Our method of RNA extraction was adapted and modified based on the protocol previously reported by Kelly et al. 2014 ^(40)^. RNA isolation from peri-implant tissue was performed using Trizol (Life Technologies, Carlsbad, CA) and RNeasy Mini kit (Qiagen, Germantown, MD). Briefly, once the samples were removed from −80°C, 1 mL of cold Trizol was immediately added to each sample. Then, the samples were homogenized (TissueLyser II, Qiagen, Germantown, MD) for 3 minutes at 30 Hz. 200 μL of chloroform was added to the samples and vortexed for 15 seconds. Next, the samples were centrifuged for 15 minutes at 4°C (13,000 rpm). The aqueous phase, approximately 600 μL, was removed and added to 600 μL of 70% ethanol to precipitate nucleic acids. RNA purification was performed using an RNeasy Mini kit (Qiagen) according to the manufacturer’s instructions, including DNase digestion (RNase-free DNase kit, Qiagen). A final volume of 25 μL was eluted.

RNA purity and quantity were determined using a spectrophotometer (Nanodrop One, Thermo Fisher Scientific, Waltham, MA). In addition, RNA Integrity Number (RIN) and quality parameters were assessed (2100 Bioanalyzer Instrument, Agilent Technologies, Santa Clara, CA). RNA integrity numbers were 8.3±0.7 and 7.6±0.6 for young and aged mice, respectively. RNA samples with low RIN and quality were discarded from the study.

### RNA sequencing

TruSeq RNA Sample Preparation Kits v2 were utilized for cDNA library preparation (Illumina, San Diego, CA) ^(41)^. Briefly, poly-A containing mRNA was isolated, converted into cDNA, and end-repaired and ligated to sequencing adapters. The resultant products were column-purified and PCR-enriched to generate final Illumina-compatible libraries (Agencourt AMPure XP, Beckman Coulter, Brea, CA). The libraries from young peri-implant tissues (n=6) and aged peri-implant tissues (n=5) were sequenced by Weill Cornell Medicine Epigenomics Core Facility using a HiSeq2500 with 50-bp single-end reads to a depth of ~15 - 25 million reads per sample.

### RNA-seq analyses

Read quality assessment and adapter trimming were performed using fastp. Reads that passed quality control filtering were then mapped to the mouse genome (mm10) and reads-in-exons were counted against Gencode v27 annotation with STAR aligner ^(42,43)^. Differential gene expression analysis was performed with edgeR using the quasi-likelihood framework in R environment.

Genes with low expression levels (<3 counts per million in at least one group) were filtered from all downstream analyses. The Benjamin-Hochberg false discovery rate (FDR) procedure was used to correct for multiple testing ^(44)^. Genes with FDR (corrected P-value) less than 0.01 and log2 fold change greater than 1 were considered differentially expressed. Gene expression changes were evaluated for enrichment of known signaling pathways with the Quantitative Set Analysis for Gene Expression (QuSAGE) method. Data visualization and downstream analyses were performed in R using a Shiny-driven visualization platform (RNAseq DRaMA) developed at the David Z. Rosensweig Genomics Research Center at the Hospital for Special Surgery (https://gitlab.com/hssgenomics/Shiny).

### RNA-seq cell type decomposition

To perform bulk RNA-seq cell type decomposition, we used the SCDC algorithm that utilizes cell-type specific gene expression information from single-cell RNA sequencing (scRNA-seq) experiments ^(45)^. To generate cell-type specific scRNA-seq reference data sets, we extracted raw scRNA count matrices from the Panglao database of scRNA-seq data that covered 2,046,548 cells from 969 experiments/samples that belonged to 119 cell types in mice ^(46)^. To improve the quality of prediction and decrease the search space, we performed rigorous quality filtering at both the experiment and individual cell levels. We first removed all cells with no assigned cell type. We then removed all individual cells with the total percentage of mitochondrial counts > 5% for cell types found in multiple experiments. We focused only on polyadenylated RNAs and selected 33,979 genes. After filtering, we created a database of 1,837,691 high quality cells representing 855 experiments and 119 cell types. Finally, we limited the search space to 37 cell types that we expected to encounter in a typical bone-related surgical material (Supplemental Figure 1). We refer this assemblage as bone-immune cell set (BICS). One hundred cells were randomly sampled for each cell type from BICS; for seven cell types in BICS that were represented by fewer than 100 cells, we used all cells available. The within-cell-type randomized reference sets were used by SCDC to determine the cell-type proportion for each bulk RNA-seq replicas (2 groups, 5 replicas for aged and 6 for young mice). The reference resampling was performed 100 times and the median cell type proportion was determined for each bulk RNA-seq replica and for both young and aged samples overall.

### Immunofluorescence staining of peri-implant tissue

One-week post-implantation, right tibiae were rapidly dissected, and soft tissue, fibulae, and malleoli were removed. Implants with attached tissue (peri-implant tissue) were removed from the tibia (n = 3 per group) (Figure 1). Then, the implants were polished using Grit 320/P400 sandpaper to remove one end of the tibial plateau. The implants were fixed with 4% paraformaldehyde (PFA) for 1 hour. After 4% PFA was removed, the implants were washed with 1XPBS for 5 minutes and permeabilized with 0.3% Triton X-100 for 15 minutes at room temperature, respectively. Next, the implants were washed once with 1XPBS for 5 minutes and blocked with 5% donkey serum for 1 hour at room temperature. Subsequently, the primary antibody, CD3 (NBP2-43674, Novusbio), was separately diluted in 5% donkey serum (1:100) and incubated for overnight at 4°C. After the removal of primary antibody, secondary antibody, Alexa fluor plus 555 donkey anti-mouse IgG secondary antibody (A32773, Invitrogen) was diluted in 1XPBS (1:400) and applied for 1 hour at room temperature. Then, the implants were washed once with 1XPBS and incubated in DAPI for 10 minutes at room temperature. Then, the implants were washed once with 1XPBS and placed in 8-well chamber slides (Thermo Fisher Scientific, Waltham, MA) (Figure 1). The images were collected as z-stacks at 10x magnification using confocal microscopy (Zeiss LSM 880).

## RESULTS

### Suppressed induction of Notch, angiogenesis and Wnt pathways at bone-implant interface tissue in aged mice

To gain insight into how aging affects osseointegration at the bone-implant interface, we performed RNA-seq of implant-associated tissue from old versus young mice one week after implantation. This time point corresponds to early osseointegration where deposition of woven bone overlaps with a resolving inflammatory response ^(15,19,23,33,35,40)^.

A total of 872 genes were differentially expressed between old and young mice (FDR < 0.01, fold change > 2), out of which 696 were expressed at a higher level in aged mice than in young mice (Figure 2A, Supplemental Table 1). To assess differences between old and young peri-implant tissue, we performed pathway enrichment analysis using QuSAGE with the MSigDb c2 set of curated pathways. Interestingly, among pathways with elevated gene expression in young mice, we noticed several bone anabolic pathways including Notch (Figure 2B, pathways 9,10, and 12, Supplemental Table 2) and Wnt-related pathways (Figure 2B, pathway 11). Further inspection of differentially expressed genes revealed that multiple genes in the Notch pathway, including Notch ligands (*Jag1, Jag2, Dll1, and Dll4*), receptors (*Notch3*, *Notch 4*), and downstream transcription factors (*Hey1, Hey2, Heyl, and Hes1*) were more highly expressed in young peri-implant tissue (Figure 2A, upper right quadrant, orange dots and Figure 2C). Similarly, several Wnt ligand receptors (*Fzd4, 5, 8, and 9*) and Wnt-regulated transcription factors (*Tcf7l1, Tcf7l2 and Tle2*) were elevated in young mice (Figure 2A and 2C). Regarding expression of transcription factors that are important in osteoblast differentiation, *Sp7* (which encodes Osterix) was not affected by aging, whereas *Runx2* was higher in young mice (Figure 2A). These results show that osseointegration at the implant surface in old mice is characterized by defective activity of the Notch pathway, which is important for both angiogenesis and expansion of bone progenitor cells. Expression of genes of the bone anabolic Wnt pathway during implant osseointegration is also compromised to a lesser extent in aged mice.

**Figure 2.**
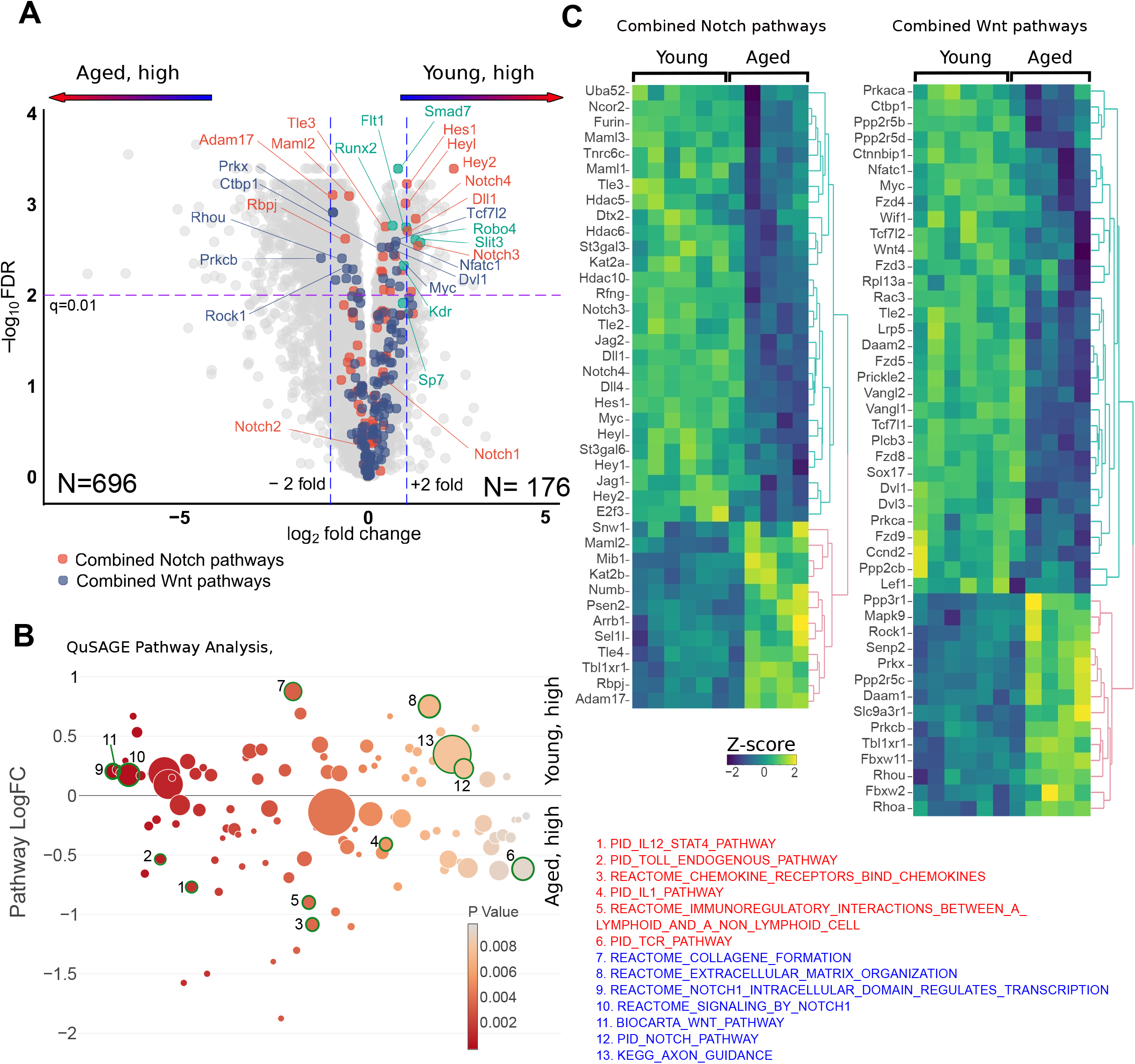
Differential expression genes involved in bone metabolism and maintenance in the peri-implant tissue from young and aged mice. **A.** A volcano plot for gene differential expression analysis of peri-implant tissue from young (n= 6) and aged (n= 5) mice. Color highlight genes from the combined Notch– and WNT-related pathways indicated at the bottom. Genes involved in angiogenesis are shown in green. **B.** QuSAGE differential pathway analysis of the peri-implant tissues from young and aged mice. All pathways with less than 10 genes were discarded and only pathways with p-value <0.01 are shown. The Y-axis shows pathway-wide log-transformed fold change between young and aged mice and the X-axis shows the p-value. The size of each circle is proportional to the number of genes in a pathway. Pathways that are upregulated in aged mice (red) and young mice (blue) are listed. **C.** Heat maps of standardized log-transformed cpm for differentially expressed genes from the Notch and WNT pathways. The columns represent individual samples from young and aged mice. Clustering of genes was performed using complete linkage method with the Euclidean distance.

Angiogenesis is a process of new blood vessel formation that is promoted by VEGF and Notch signaling ^(47–49)^ and is important for new bone formation, fracture healing, and osseointegration ^(15,50–53)^. Recent work has implicated specialized Endomucin-high endothelium, which is regulated by the SLIT3/ROBO1 pathway, in bone vascularization and titanium implant integration ^(15,54)^. Treatment with recombinant SLIT3 improves bone fracture healing and prevents bone loss ^(55–57)^. Importantly, we observed that both *Slit3*, which appears to be osteoblastic specific ^(57)^, and *Robo4*, which can be considered as an endothelial maker ^(57)^, were more highly expressed in the peri-implant tissue of young mice (Figure 2A, green dots). Similarly, two VEGF receptors, *Kdr* and *Flt1* were expressed at a higher level in young mice (Figure 2A, green dots), further supporting the notion of compromised angiogenesis in old mice.

### Increased activation of immune and inflammatory pathways at the bone-implant interface in aged mice

Strikingly, the expression of multiple immune and inflammatory pathways was significantly increased in old relative to young mice including IL-12-STAT4 signaling (Figure 2B, pathway 1), innate immune activation (Toll-like receptor (TLR) and IL-1, pathways 2 and 4), and cytokine and chemokine signaling (pathway 3,5, and 6). Activation of both innate and adaptive immune components was supported by a significant elevation of TLR signaling (pathway 2) and a group of pathways related to T cell receptor (TCR) signaling (e.g. pathway 6). Toll like receptors are potent activators of inflammatory NF-κB signaling in immune cells and are activated by tissue damage products, whereas TCR signaling is suggestive of activation of lymphocytes.

Expression of the core components of the TCR (*Cd3e*, *Cd3d*, *Cd3g*), co-receptors *Cd4*, *Cd28*, and costimulatory molecules *Cd80/Cd86* was elevated in aged mice (Figure 3A, top panel and Figure 3B). As expression of these molecules typically does not change with cell activation, this result is suggestive of increased infiltration of T cells, including CD4+ T cells, into peri-implant tissue in old mice. Aged peri-implant tissues also exhibited increased expression of *Stat4*, which is induced during differentiation and activation of CD4+ T helper 1 (Th1) cells, and of *Il12*, a major inducer of Th1 cells. Activated Th1 cells produce IFN-γ and TNF; and, accordingly, expression of *Tnf* and canonical IFN-γ target genes like *Ccl5* was elevated in aged peri-implant tissues (Figure 3A, middle panel).

**Figure 3.**
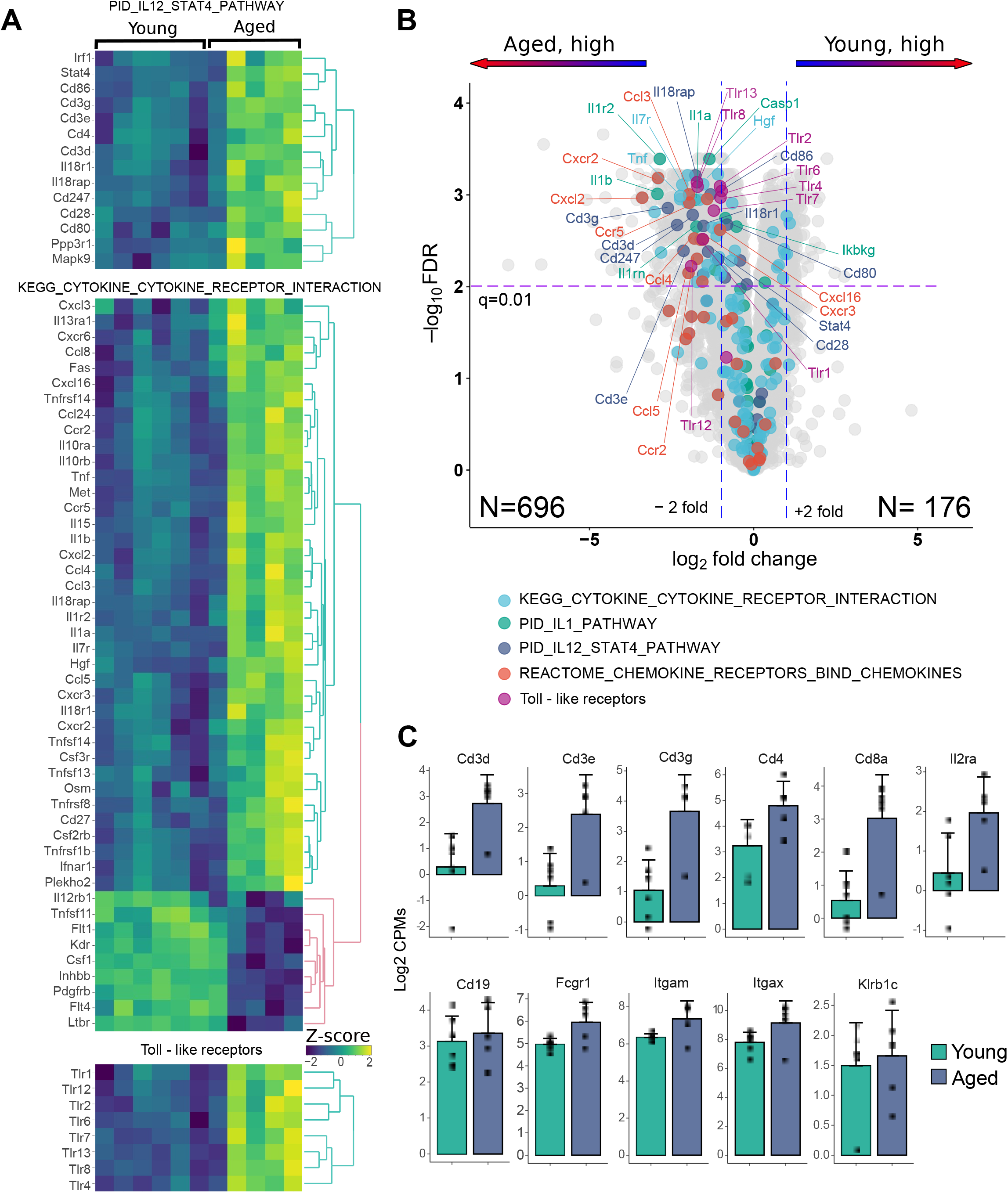
Differential expression inflammatory genes in the peri-implant tissue from young and aged mice. **A.** Heat maps of standardized log-transformed cpm of differentially expressed immune-inflammatory genes from the IL12_STAT. The columns represent individual samples from young and aged mice. Clustering of genes was performed using complete linkage method with the Euclidean distance. **B.** A volcano plot for gene differential expression analysis of peri-implant tissue from young (n=6) and aged (n=5) mice. Color highlight genes from the pathways indicated at the bottom of each panel. **C.** Comparison of cell surface marker expression in young and aged maps: T cells - Cd3d, Cd3e, Cd3g, Cd4, Cd8a, Il2ra(Cd25), B cells – Cd19, Monocytes/Macrophages Fcgr1 (Cd64), Itgam (Cd11b), dendritic cells – Itgax (Cd11c), NK cells – Klrb1c (NK1.1).

Major mediators of innate immunity include the cytokines IL-1, TNF, and TLRs which activate inflammatory genes via NF-κB signaling. IL-1 is expressed predominantly by innate myeloid cells such as macrophages and dendritic cells, which are also major producers of TNF. The bone-implant interface tissue in aged mice exhibited increased expression of *Il1a*, *Il1b*, and *Tnf* and various downstream inflammatory NF-κB target genes and chemokines such as *Cxcl2*, *Ccl3*, *Ccl4*, and *Ccl8*. Thus, an innate immune response is more strongly activated in aged than young mice in response to bone injury and tibial implantation. TLRs are highly expressed on various immune cells and their expression can increase during cell activation. Strikingly, expression of 8 out of 13 known TLRs was elevated in aged relative to young mice (Figure 3A, bottom panel). This includes TLRs that can sense tissue degradation products such as collagen and fibronectin fragments (TLR2, TLR4) and nucleic acids (TLR7/8) and chromatin proteins (TLR2) released by dying cells ^(58)^.

We then surveyed the relative expression of additional canonical immune cell markers in bone-implant interface tissue (Figure 3C). In addition to *Cd3d* and *Cd4*, expression of the CD8+ T cell marker *Cd8a* and of macrophage and DC marker genes *Itgam*, *Itgax* and *Fcgr1* was increased in aged peri-implant tissue, whereas B cell (*Cd19*) and NK cell (*Klrb1c*) marker gene expression was comparable in aged and young tissues.

To address more thoroughly how increased expression of inflammatory mediators in aged mice may reflect increased infiltration of immune cells, we performed cell-type decomposition of bulk RNA-seq data using SCDC deconvolution method ^(45)^. SCDC algorithm uses the cell-type specific gene expression reference sets generated from multiple single-cell RNA-seq experiments (see Materials and Methods). Comparison of cell type composition in peri-implant tissues revealed substantial differences between young and aged mice. In young mice one week after surgery, cells expressing osteoblast lineage genes represent the largest proportion of cells in the bulk RNA-seq samples (>50%); this is in accord with mobilization of stromal precursors which differentiate along the osteoblast pathway to lay down bone. Fibroblasts and macrophages were the distant next two common cell types in young peri-implant tissue (Figure 4A). In aged mice, the proportion of osteoblast-lineage cells contributing to bulk RNA-seq gene expression was decreased considerably (24%), and the macrophages became the largest contributing cell type (28%). Concomitantly, several other immune cell types were elevated in old compared to young mice (Figure 4B, Supplemental Figure 1), although these cell types were present at a much lower level.

**Figure 4.**
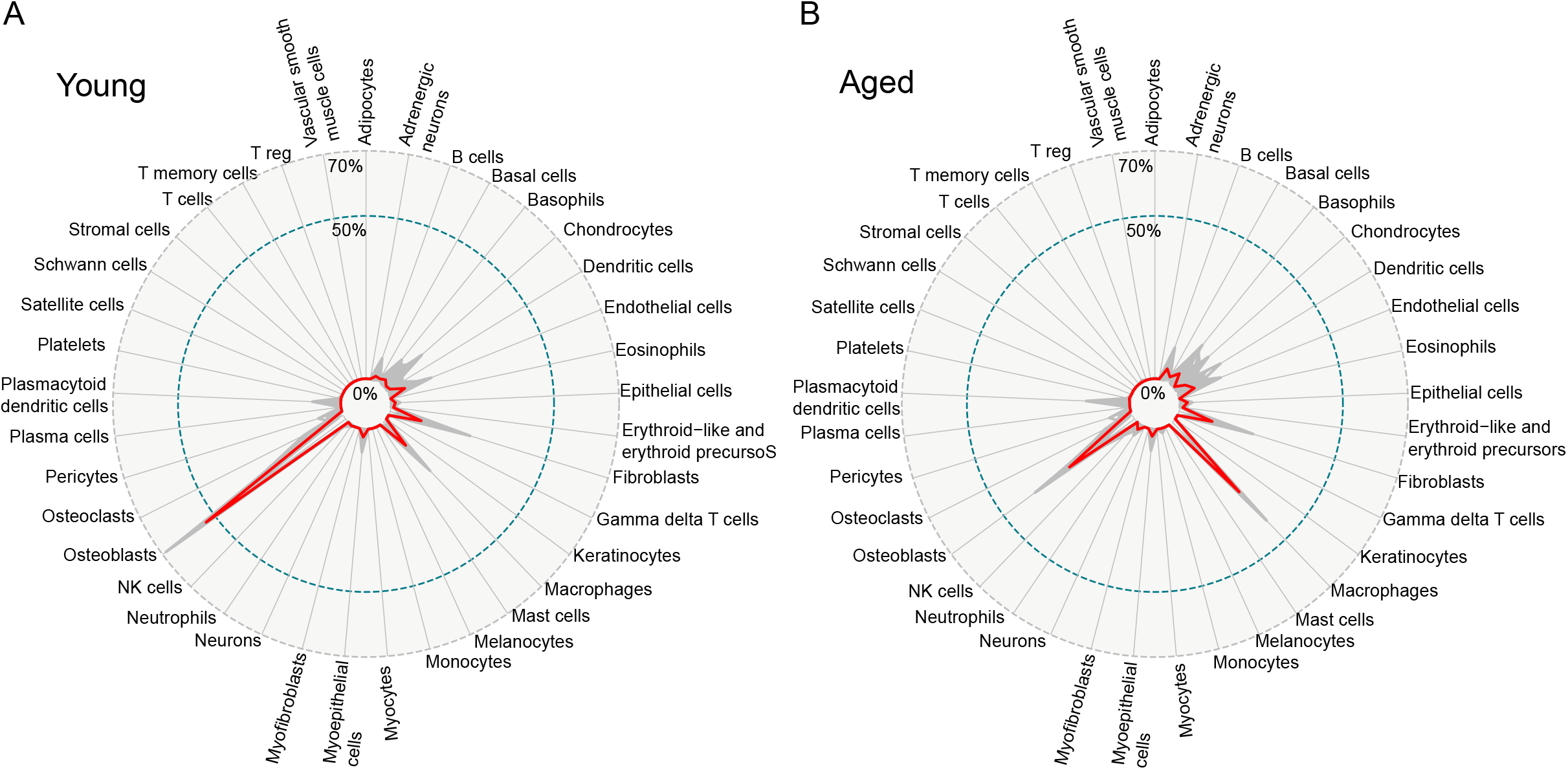
Cell type decomposition of bulk RNA-seq data from young (A) and aged (B) peri-implant tissue. Cell type composition has been inferred as described in the Materials and Methods section. The red line represents the median proportion of each cell type from BICS for all bulk RNA-seq replicas and all 100 bulk-to-scRNA-seq reference comparisons. The thin gray lines represent the average proportions of each group (young versus aged) for each cell type from BICS across all the iterations to show the variation in each SCDC proportional output.

Overall, these findings suggest that bone injury and implant insertion elicit increased immune cell infiltration and stronger innate immune response at the bone-implant interface in aged mice relative to young mice.

Similarly, we have additionally performed a differential expression and pathway analyses in cancellous bone adjacent to the implant in aged versus young mice (Supplemental Figure 2A). We noted a highly significant difference in transcripts associated with bile acid and bile salt metabolism pathway (Supplemental Figure 2A, pathway 1, Supplemental Table 3). This was primarily due to elevated expression of cholesterol 25-hydroxylase (*Ch25h*) (Supplemental Figure 2B) in aged mice, consistent with age-related differences in cell metabolism. Large differences were also observed in the expression of genes involved in protein secretion and matrisome associated gene sets (pathways 3 and 4), which combine many proteins of extracellular matrix, ECM-modifying enzymes and soluble growth factors ^(59)^. This finding likely reflects basal differences in bone metabolism and response to injury between young and aged mice.

### T cell infiltration at the bone-implant interface

After the observation of expression of T cell marker and activation genes (Figure 3A-C) in aged mice, we wished to confirm the presence of T cells at the bone-implant interface using immunofluorescence microscopy. This is technically challenging as the peri-implant tissue remains attached to the titanium implant after removal of the implant from the tibia. Thus, we developed an approach using confocal microscopy for 3D immunofluorescence imaging of tissue adherent to the implant surface. CD3 staining was readily observed in implant-associated tissue, supporting infiltration of T cells into peri-implant tissue (n=3/group, Figure 5).

**Figure 5.**
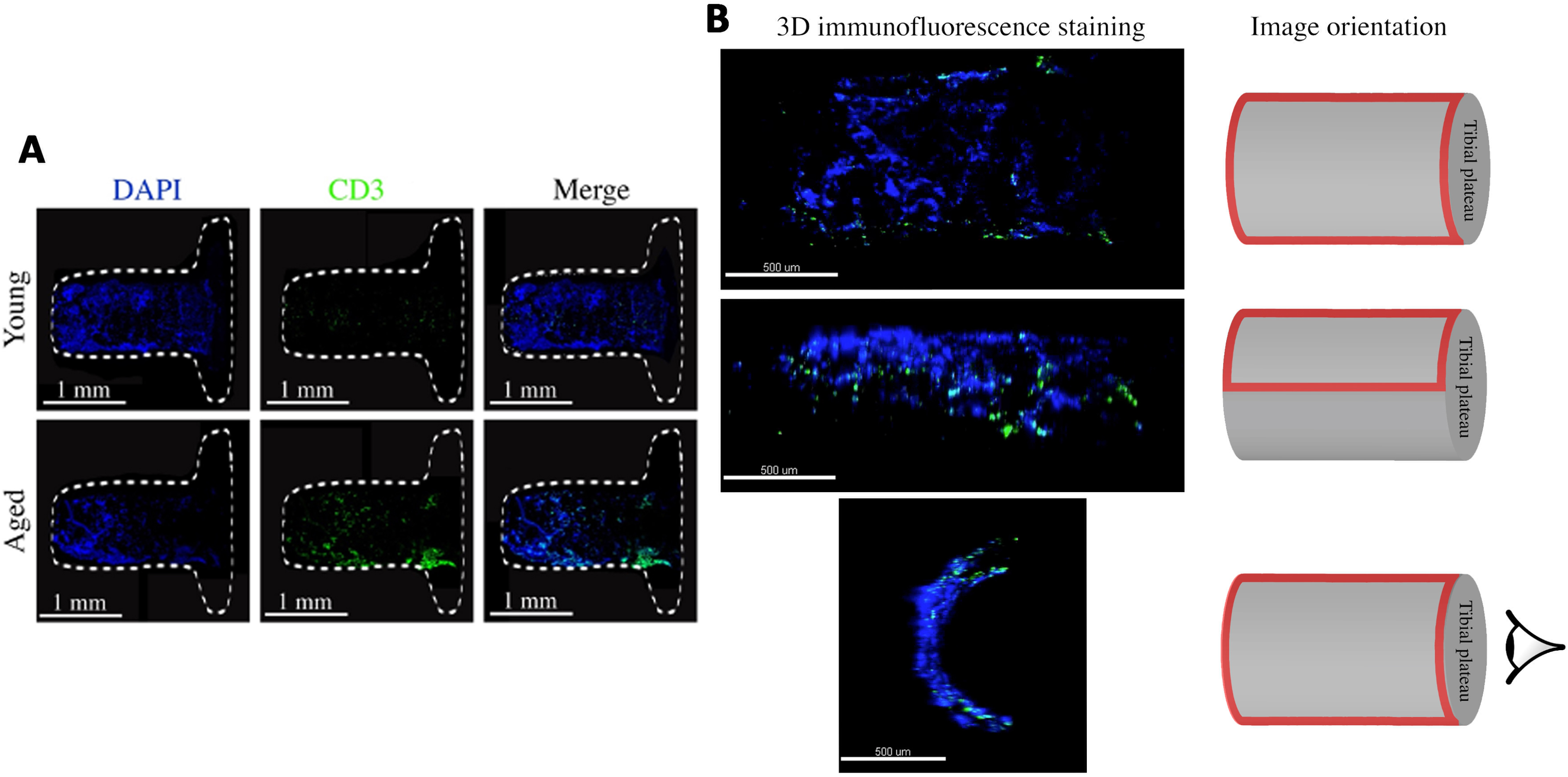
3D Immunofluorescence staining of the implant surface harvested from young and aged mice. **A.** After one-week post-implantation, the implants were polished, fixed, and stained using alexa fluor plus 555 donkey anti-mouse IgG secondary antibody to detect CD3 (green) and DAPI to label nuclei (blue) (10x, z stacks). The white dash lines represent the outline of the implant. **B.** Confocal imaging and image orientation. The red frames indicate the image region.

## DISCUSSION

The mechanistic basis for poor osseointegration in elderly patients is not understood, but increased inflammation that occurs with aging (‘inflamm-aging’) has been implicated in tissue degeneration and poor healing in other systems ^(60–63)^. Here, we have extended our clinically relevant, mechanically loaded murine model of TJR through the development of new methods to address the technical challenge of performing transcriptional profiling of the very limited peri-implant bone tissue. Our results reveal age-related decreases in angiogenic and bone anabolic pathways associated with increased immune activation at the site where injured bone integrates with a clinically relevant porous titanium implant. These findings highlight the importance of augmenting angiogenic and bone anabolic pathways in elderly patients undergoing TJR, and suggest immune modulation as a novel therapeutic strategy to enhance implant osseointegration.

We performed our studies one week post-implantation, when the initial inflammatory response to injury is subsiding and the bone formation phase is underway ^(15,19,23,33,35)^. At this phase, increased and sustained inflammation in aged mice mediated by innate immune cytokines such as TNF and IL-1 can suppress osseointegration by augmenting osteoclastogenesis, suppressing osteoblasts directly or via increased production of Wnt pathway inhibitors such as DKK1 ^(32,64)^, and compromising the emergence of a pre-resolution immune response. Innate immune cells are directly activated by tissue damage and degradation products, activate tissue resident cells, and induce production of chemokines that attract additional innate and acquired immune cells from circulation to the site of injury. We unexpectedly observed increased T cell infiltration and an IFN-γ gene signature in the peri-implant tissue, indicative of IFN-γ production. It is not yet clear whether IFN-γ at the site of osseointegration is produced by CD4+, CD8+, and γδ T cells or NK cells, and how these cells become activated. However, given its role in suppressing fracture repair, bone formation, Notch signaling, and angiogenesis, it is likely IFN-γ contributes to impaired osseointegration in aged mice ^(35,36,65,66)^.

Our data demonstrate that osteogenesis, angiogenesis, and inflammation occur directly at the peri-implant site within one week after implantation. Aged mice exhibit impaired expression of genes associated with vascular formation and preferential suppression of Notch signaling relative to other anabolic pathways (Wnt, BMP, PTH), suggesting that Notch-mediated osteogenic mechanisms are disproportionately affected by aging. Diminished Notch pathway activity may contribute to decreased osseointegration by decreasing angiogenesis in addition to decreased endochondral and intramembranous ossification ^(67,68)^. The notion that elevated immune and inflammatory responses in elderly organisms may suppress bone anabolic, or promote resorptive, pathways is consistent with previous reports of decreased fracture healing with age ^(37,69–71)^.

Typically, histological evaluation of clinical implant samples is performed after patients experience implant failure. The absence of samples from early stage osseointregation failure and successful osseointegration impair our ability to understand the early steps in ossteointegration failure Furthermore, the existence of metal implant in the bone makes sample preparation difficult. Therefore, using our murine model, we developed a novel technique of 3D immunofluorescence staining to analyze the peri-implant (surface) tissue without cutting our specimens. This implant surface staining technique can be utilized to monitor the process of bone formation on the implant surface or identify presence of bacteria in a bone infection animal model. In addition, the approach can be applied to evaluate bone healing on the surface of various implant materials.

Our study has several limitations. The gene expression data analysis was based on mixed cell populations. This approach identifies changes in cell composition based on marker gene expression and cell deconvolution algorithms, but it is more difficult to assess cell activation states and which cell populations are being activated. In addition, this study only evaluated one time point, which does not reflect the entire process of bone healing after implantation. Future studies should focus on finding methods to enhance osteogenesis and angiogenesis and regulate inflammation in elderly patients. Our model can provide a viable platform for preclinical studies. Furthermore, this study opens the possibility that targeting the Notch pathway, promoting angiogenesis, or modulating the immune response at the peri-implant site can enhance osseointegration and improve the outcome of joint replacements in older patients.

## ACKNOWLEDGMENTS

We would like to acknowledge the Weill Cornell Medicine Microscopy and Image Analysis Core Facility for image analysis consulting, the Weill Cornell Medicine Genomics Resources Core Facility and Weill Cornell Medicine Epigenomics Core Facility for nucleic acid quality assessment and RNA sequencing, and the David Z. Rosensweig Genomics Research Center at the Hospital for Special Surgery for RNA sequencing data analyses. MBG holds a Career Award for Medical Scientists from the Burroughs Wellcome Fund, awards from the NIH under DP5OD021351 and R01AR075585 and an award from the Pershing Square Sohn Cancer Research Alliance. The Rosensweig Genomics Center is supported by The Tow Foundation. LBI was supported by NIH grants DE019420 and AI044938. XY was supported by Grant UL1 TR000457 of the Clinical and Translational Science Center at Weill Cornell Medicine and Feldstein Medical Foundation.

