## Supplementary figures and images for "Age-related Impairment of Implant Osseointegration is Associated with Immune Activation and Suppression of Angiogenic, Notch, and Wnt Pathways"

### Supplemental Figure 1

Supplemental figure 1

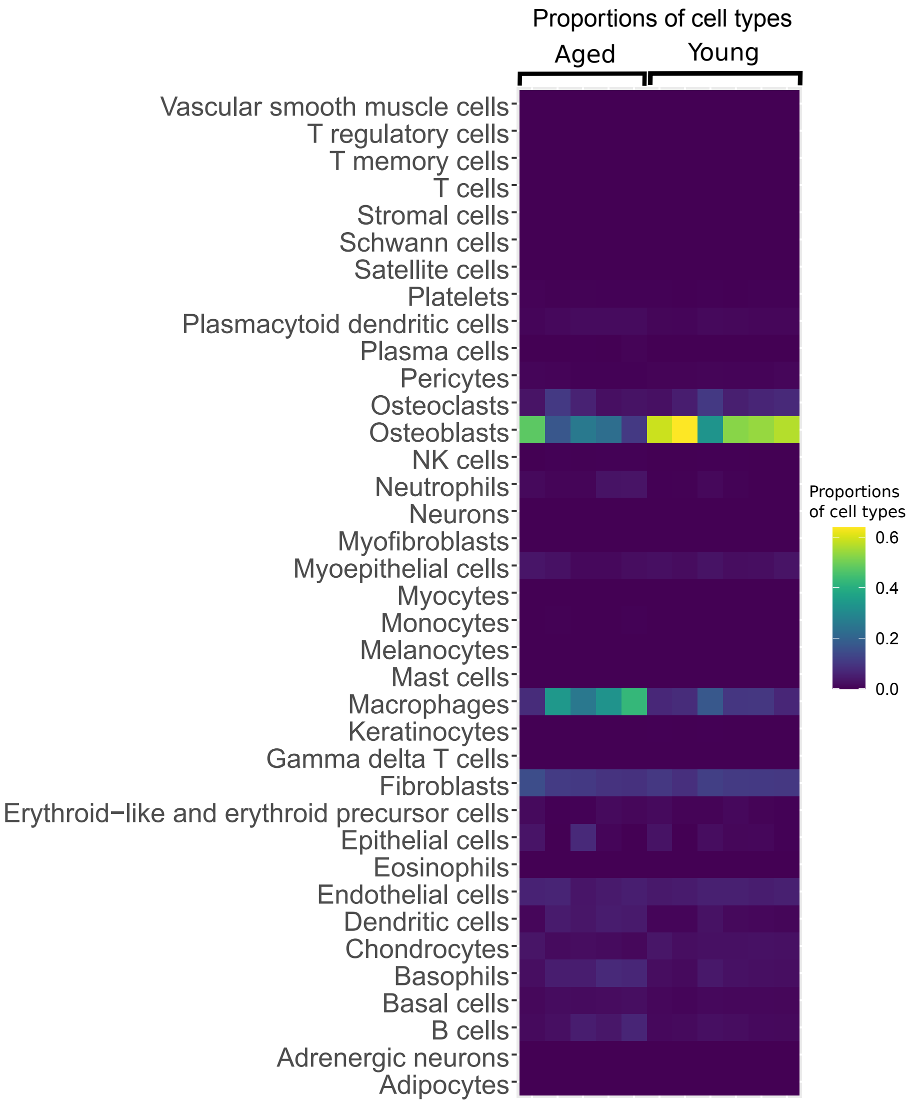

### Supplemental Figure 2

Supplemental figure 2

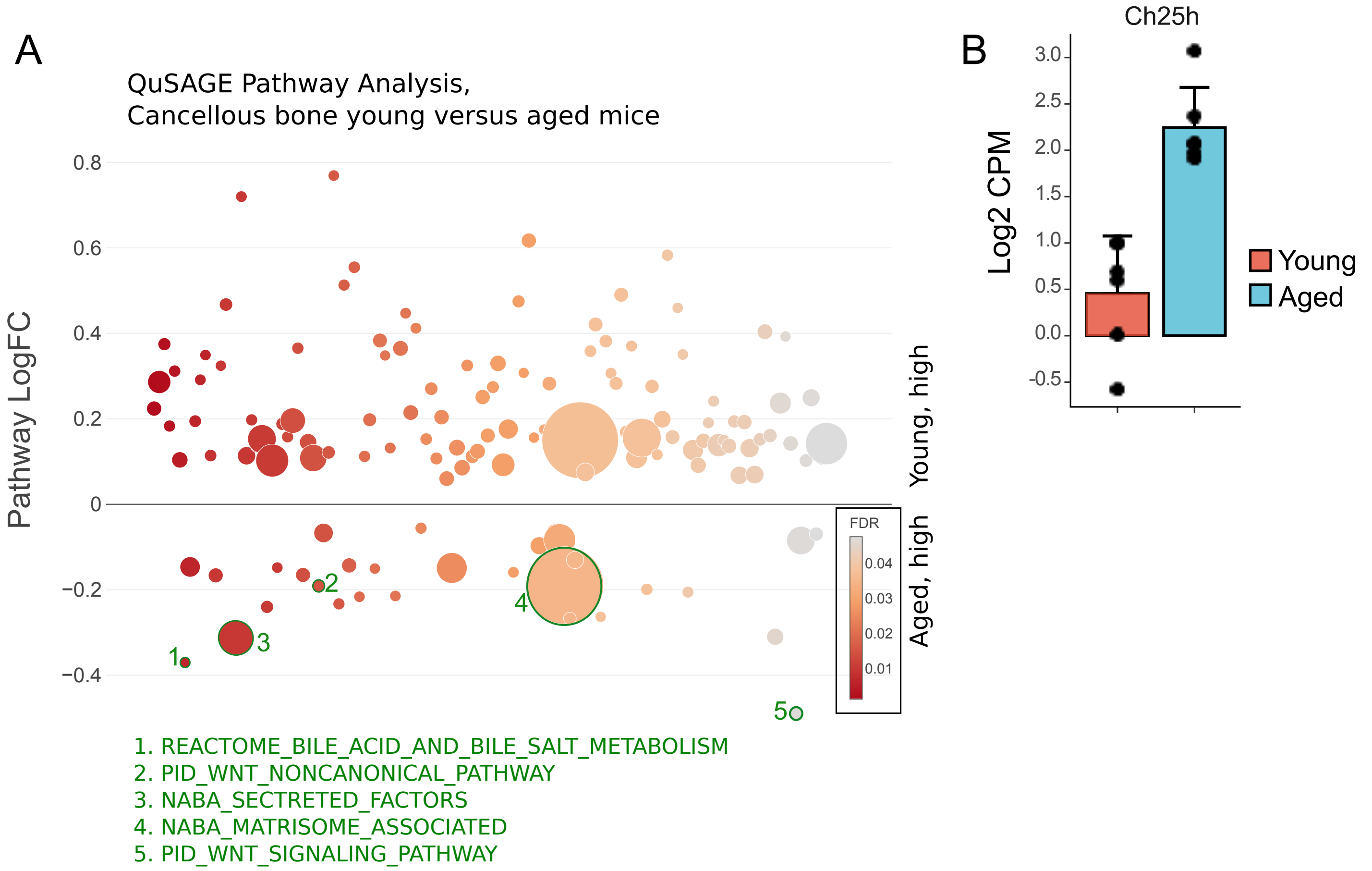
